# Predicting climate-driven distribution shifts in *Hyalomma marginatum* (Ixodidae)

**DOI:** 10.1101/2023.04.01.535198

**Authors:** Olcay Hekimoglu, Can Elverici, Arda Cem Kuyucu

**Affiliations:** Biology Department, Hacettepe University, Ankara, Turkey; Biodiversity Institute, University of Kansas, Lawrence, Kansas, United States

**Keywords:** *Hyalomma marginatum*, Ticks, Ecological Niche Modelling, Maxent, Climate change, GCMs, Mediterranean Basin, Europe, Anatolia, CCHF

## Abstract

*Hyalomma marginatum* is an important tick species which is the main vector of Crimean-Congo haemorrhagic fever (CCHF) and spotted fever. The species is predominantly distributed in parts of Southern Europe, North Africa and West Asia. However, due to ongoing climate change and increasing reports of *H. marginatum* in central and northern Europe, the expansion of this range poses a potential future risk. In this study, we employed an Ecological Niche Modelling (ENM) approach to model the current and future climatic suitability of *H. marginatum*. Using high resolution climatic variables from the Chelsa dataset and an updated list of locations for *H. marginatum*, we constructed 2 models under current environmental conditions using MaxEnt: a more specific model for only current conditions, and a more general model for both current conditions and future projections under the ssp370 and ssp585 scenarios. Our models show that the climatically suitable region for *H. marginatum* matches the current distributional area in the Mediterranean basin and West Asia. When applied to future projections, our models suggest a considerable expansion of *H. marginatum*’s range in the north in Europe as a result of rising temperatures. However, they also predict a decline in central Anatolia, potentially due to the exacerbation of drought conditions in that region.

**Subjects:** Ecology, Entomology, Epidemiology, Infectious Diseases, Climate Change Biology

## INTRODUCTION

Human intervention in ecosystems, including globalization, urbanization, rapid transportation, land use, and climate change, has led to a global increase in the transmission rate of several pathogens and vector organisms that serve as reservoirs for these pathogens (Aguilar-Domínguez et al., 2021; Andersen & Davis, 2017; Carvalho et al., 2017; Kovats et al., 2001; Leder et al., 2021; Semenza & Suk, 2018).

The relationship between vector populations and climate has gained prominence as the effects of anthropogenic climate change become more apparent, with increases in mean annual temperatures and a dramatic rise in the frequency and magnitude of extreme weather events (Harris et al., 2018; Mckechnie & Wolf, 2010; P. Stott, 2016; P. A. Stott et al., 2016). While the most severe impacts of climate change are expected to affect ectotherm populations in the tropics, population growth rates of insects and other arthropods are also anticipated to increase in mid-to-high latitude regions due to the influence of warmer temperatures on growth rates (Bonebrake & Deutsch, 2012; Deutsch et al., 2008; Rocklöv & Dubrow, 2020). In addition to the effects of temperature on the biology of disease transmitting vector organisms, increased precipitation and extreme precipitation events in some regions may lead to an increase in vectors, whereas longer dry periods could increase tick mortality (Githeko et al., 2000; Rocklöv & Dubrow, 2020).

Ticks are the vectors of several critical pathogens and pose a significant threat to animal and public health. The genus *Hyalomma* Koch, 1844 is of a particular concern due to its role as the vector of Crimean Congo Hemorrhagic Fever (CCHF), babesiosis and Rickettsia in the Mediterranean region, Africa and Asia (Duscher et al., 2018; Ros-García et al., 2011; Sultankulova et al., 2022; Vatansever et al., 2007). Specifically, *Hyalomma marginatum* Koch, 1844 presents a substantial health threat as the main vector of CCHF (Bonnet et al., 2022; Ergönül, 2006; Gale et al., 2010). Effective and sustainable control of ticks, like many arthropod pests and vectors, can be achieved with a solid understanding of the species’ biology, ecology and distribution (Bonnet et al., 2022). Knowledge of *H. marginatum*’s current distribution is crucial for projecting areas at risk for CCHF expansion. Recent reports have shown that the vector of CCHF, *H. marginatum* Koch, 1844 has expanded its geographic range and shifted northward into previously unoccupied areas (Bah et al., 2022; McGinley et al., 2021; Vial et al., 2016). Furthermore, the expansion of the suitable areas has been documented in Mediterranean countries, central Europe and Balkans (Fernández-Ruiz & Estrada-Peña, 2021). Recent studies have highlighted that northern latitudes are becoming more suitable for *H. marginatum* activity and survival as a result of climate change (Chitimia-Dobler et al., 2019; Grandi et al., 2020; McGinley et al., 2021). The Mediterranean region is among the areas projected to be most negatively affected by climate change (Bardsley & Edwards-Jones, 2007; Newbold et al., 2020; Ulbrich et al., 2006). Studies have indicated an increase in CCHF transmission in the Mediterranean basin (Maltezou & Papa, 2010) and a tendency for the northward expansion of CCHF’s range in the Mediterranean (Andersen & Davis, 2017; Estrada-Peña & Venzal, 2007; Fernández-Ruiz & Estrada-Peña, 2021; Williams et al., 2015).

Thanks to advances in machine learning algorithms in ecology, such as MaxEnt (Crisci et al., 2012; Elith et al., 2011; Merow et al., 2013; Phillips & Schapire, 2004) and the advent of high-resolution climatic datasets such as Chelsa (Karger et al., 2017) and WorldClim (Fick & Hijmans, 2017), ecological niche modeling (ENM) has become a fundamental method for investigating possible current and future distributions of disease-transmitting vectors over the past two decades (Aguilar-Domínguez et al., 2021; Alkishe et al., 2021; Chalghaf et al., 2018; Moo-Llanes et al., 2021; Raghavan et al., 2020). Correlative ecological niche models primarily rely on combining georeferenced species records with predictive environmental variables (e.g., climatic, topographic, soil) to build a coefficient matrix representing the organism’s multi-dimensional niche (Peterson & Soberón, 2012; Sillero & Barbosa, 2021; Warren & Seifert, 2011). ENM’s have been widely used to predict the potential future distributions of vectors under various global circulation scenarios of climate, in addition to their current potential (Aguilar-Domínguez et al., 2021; Alkishe & Peterson, 2022; Wu et al., 2022).

The main objective of this study is to predict climatically suitable areas for *Hyalomma marginatum* under both present-day conditions and future climate scenarios. To achieve this aim, bioclimatic parameters for present and near-future scenarios (2011-2040 and 2041-2070) obtained from the CHELSA database and *H. marginatum* locations from the literature and previous field surveys by the authors were used to build ecological niche models with MaxEnt to show suitable areas for present and future projections. The findings of this study will help predict potential new suitable areas in the Mediterranean basin and Europe that *H. marginatum* may colonize in the future and enhance surveillance efforts in areas identified as high risk.

## MATERIAL & METHODS

### Occurrence Records

Local data from Turkey were derived from published literature up to 2021, which used both morphological and molecular methods to identify tick samples (Hekimoglu et al., 2020; Hekimoglu & Ozer, 2017) and field collection data from 2018 (n = 13 locations) and 2021 (n= 7 locations). Morphological identification was conducted using taxonomic keys (Apanaskevich & Horak, 2008; Estrada-Peña et al., 2018). Since *H. marginatum* populations show morphological variations (Apanaskevich & Horak, 2008) and *Hyalomma* species have been one of the most misidentified taxa using morphological methods (Estrada-Pena & De La Fuente, 2016), conducting a molecular step is extremely important. For this purpose, mitochondrial 16S rDNA sequences (Mangold et al., 1998) were generated for samples of localities where molecular data were not generate before. In total, 66 geographical points from Turkey were included in the analyses.

To represent the distribution area of the species worldwide, we included geographic locations of

*H. marginatum* from different parts of the world from a previous compilation by Estrada-Pena & de La Fuente (2016), composed of literature reviews between 1970 and 2014. Firstly, the geographic information of *H. marginatum* was recorded on a separate data sheet. Secondly, these raw records were cleaned and reduced by removing localities, (1) where tick collection had been conducted from birds since finding ticks on birds does not mean that ticks can establish populations in these areas (Estrada-Peña et al., 2011; Fernández-Ruiz & Estrada-Peña, 2021); (2) sampling information from Cyprus was removed as molecular data does not confirm that *H. marginatum* exists in Cyprus (Hekimoglu & Ozer, 2017). After these steps, a total of 565 geographic coordinates were obtained. The coordinates used in this study are available at https://data.mendeley.com/datasets/n8b3kps7nk/1.

### Environmental variables

We used Chelsa V2.1 dataset (Karger et al., 2017, 2022) available at https://chelsa-climate.org, a relatively new high-resolution (30 arc sec, ∼1km) climate dataset that includes additional important microclimatic variables in addition to the counterparts (Chelsa-Bioclim) of popular WorldClim bioclim variables (Fick & Hijmans, 2017; Hijmans et al., 2005). These include variables related with microclimate, water content of air, and humidity which are important for

*H. marginatum* (Estrada-Peña, 2023; Estrada-Peña et al., 2011). Furthermore, while the WorldClim dataset used for current distribution predictions uses extrapolations made between 1970-2000, the Chelsa dataset uses a dataset made for the period between 1980-2010. The additional important variables (Chelsa-Bioclim+) included near surface relative humidity (hurs), vapour pressure deficit (vpd), climate moisture index (cmi), growth degree days above 0C, net primary productivity (npp) and surface downwelling shortwave radiation (rsds). Also included were growth season temperature (gst) and growth season precipitation (gsp), which are indicative of *H. marginatum*, whose activity coincides greatly with the growth season (Estrada-Peña et al., 2011). Bioclim variables 8, 9, 18 and 19 from the Worldclim dataset (Fick & Hijmans, 2017) are reported to have significant spatial artefact problems visible as anomalous discontinuities between neighbouring pixels and thus it is generally advised to remove them before carrying out analyses (Aguilar-Domínguez et al., 2021; Alkishe & Peterson, 2022; Escobar, 2020; Escobar et al., 2014). We removed these variables from the Chelsa dataset since we observed that the same artefacts are also present in Chelsa-Bioclim data. All remaining variable rasters were clipped to the area of interest, which is between latitudes -20°, 60° and longitudes 20°, 60° WGS84. All geographic computations were carried out with QGIS (*QGIS Geographic Information System*, 2022) GDAL library in Python (Open Source Geospatial Foundation, 2022) and R version 4.2.2 (R Core Team, 2022).

### Future Projections

For future predictions, environmental variables from 5 different global circulation models (GCM) of ssp370 and ssp585 scenarios were used. These include UKESM1-0-LL (Sellar et al., 2019), MRI-ESM2-0 (Oshima et al., 2020), MPI-ESM1-2 (Mauritsen et al., 2019), IPSL-CM6-LR (Boucher et al., 2020) and GFDL-ESM4 (Dunne et al., 2020) for the Chelsa dataset, also available at https://chelsa-climate.org.

### Ecological Niche Modeling (ENM)

Before building the models, we removed occurrence records that fell outside environmental raster pixels. In order to reduce spatial auto-correlation, the locations for *H. marginatum* were thinned to a 5 km radius using the spThin package for R (Aiello-Lammens et al., 2015). To simulate the accessible **M** space (Barve et al., 2011; Soberon & Peterson, 2005) for *H*.*marginatum*, we created a buffered minimum convex polygon with a buffer area of 100 km around the occurrence records and environmental variable rasters representing the M space were clipped to this buffer area before building models. Thinning and the creation of the buffer zone were carried with ellipsenm package for R available at https://github.com/marlonecobos/ellipsenm (Cobos et al., 2020). We built 3 sets by setting correlation thresholds between variables using the vifcor function in the R package usdm (Naimi, 2017). The vifcor function selects the variables with lower vif (variance inflation factors) from correlation pairs to build the sets. The correlation thresholds were 0.9, 0.8, and 0.75, respectively. For the 1^st^ model dataset Set 2 and Set 3 were identical, so we used only Set 1 and Set 2 to build the models. For the Chelsa dataset some of the Bioclim+ variables are not available for future projections (vpd, hurs, rsds and cmi). Thus, we built 3 different sets not including these variables to build a separate 2^nd^ model (Model 2) for future projections.

ENM’s were built with Maximum Enthropy algorithm (Phillips & Schapire, 2004) using MaxEnt 3.4.4 (Steven et al., 2021) implemented in Kuenm package (Cobos et al., 2019) for R. Using the kuenm package, we built calibration models with all combinations of MaxEnt features L (linear), Q (quadratic), P (product), and H (hinge). All models were repeated with regularization multiplier parameters 0.1, 0.2, 0.3, 0.4, 0.5, 0.6, 0.7, 0.8, 0.9, 1, 2, 3, 4 and 5. The performance of all these models were evaluated primarily with partial Receiver Operating Characteristics (pROC) test and an omission threshold rate of 5% (Aguilar-Domínguez et al., 2021; Peterson & Soberón, 2012), secondary evaluations were done with Akaike Information Criterion corrected for small sample size (AICc). The eventual calibration models were selected from statistically significant, that have omission rates below 5% and with ΔAICc values below 2 (Nuñez-Penichet et al., 2021; Warren & Seifert, 2011).

After model calibrations, final models were created with the selected parameters in the resulting calibration models using all the same occurrences of training and testing data with 10 bootstrap replicates. Then, these final models were additionally evaluated with the independent locations data (n = 24) that were not used for building and selecting these models. Projections for the future GCM scenarios were done with clamping.

Map binarization was done by using the average logistic thresholds of maximum training sensitivity plus specificity of 10 replicates (Liu et al., 2013). Consensus maps of the niche models for the future projections were created by taking the average of GCMs.

## RESULTS

### Model parameters

Environmental parameters selected based on correlation thresholds and vif for variable sets used in model calibrations are shown in Table 1. In total 420 candidate models were created for the 1^st^ model (Model 1). All of 420 candidate models were statistically significant (pROC test p < 0.05); however, only 82 of these fulfilled the omission rate criteria of 5% (omission rate = 3.6%). Only 1 of these met the AICc criteria of ≤ 2 (ΔAICc = 0.90). This final selected model is a LQ (linear quadratic) model with a regularization multiplier parameter of 0.2, which uses variables included in Set 1 with a mean AUC ratio of 1.21. The final evaluation with the independent test set resulted with an omission rate of 8.3%. Although this value is larger than the 5% threshold we set for evaluations with the cross-validation sets, the independent test set had a sample size of n = 24, and 8.3% omission rate means that our final model classified 22 of 24 test locations correctly which is a pretty good result for a machine learning algorithm. The percent contribution and permutation importance of 12 predictors are shown in Table 2. Out of these, the most important parameter was cmi (climate moisture index) with a contribution of 33.1%, followed by Bio14 (precipitation amount of the driest month) with a 24.2% contribution, and rsds (surface downwelling shortwave radiation) with an 8.4% contribution. Bio6 (minimum air temperature of the coldest month) was the 4^th^ most important predictor (7% contribution).

**Table 1.** Table shows the environmental predictors used different sets in Model 1 and Model 2 with explanation of the predictors in the first column

|  | Model 1 |  | Model 2 |  |  |
| --- | --- | --- | --- | --- | --- |
|  | Set 1 | Set 2 | Set 1 | Set 2 | Set 3 |
| **Bio3** (Isothermality) | √ | √ | √ | √ | X |
| **Bio4** (temperature seasonality) | √ | X | √ | X | X |
| **Bio6** (Min. air temp. of the coldest month) | √ | X | √ | X | X |
| **Bio7** (annual range of air temperature) | √ | √ | √ | √ | √ |
| **Bio13** (precipitation amount of the wet-test month) | √ | √ | √ | √ | √ |
| **Bio14** (precipitation amount of the driest month) | √ | √ | √ | √ | X |
| **Bio15** (precipitation seasonality) | √ | √ | √ | √ | √ |
| **Gdd0** | X | X | √ | X | X |
| Growing degree days heat sum above 0°C |  |  |  |  |  |
| **Gst** (Mean temperature of the growing season) | √ | √ | √ | √ | √ |
| **Gsp** (precipitation amount of the growing season) | √ | X | √ | X | X |
| **Npp** (Net primary productivity) | √ | X | √ | X | X |
| **Rlds** (surface downwelling shortwave radiation) | √ | X | X | X | X |
| **Cmi** (climate moisture index) | √ | X | X | X | X |

**Table 2.** Percent contribution of environmental predictors to Model 1 and Model 2.

| Model 1 |  | Model 2 |  |
| --- | --- | --- | --- |
| Predictor | Percent contribution | Predictor | Percent contribution |
| Cmi | 33.1 | Bio13 | 31.3 |
| Bio14 | 24.2 | Bio14 | 19.4 |
| Rlds | 8.4 | Gdd0 | 13 |
| Bio6 | 7 | Bio6 | 8.2 |
| Bio15 | 6 | Npp | 8 |
| Bio13 | 5.7 | Bio15 | 7.5 |
| Gst | 4.2 | Bio4 | 4.9 |
| Npp | 4.1 | Bio3 | 4.5 |
| Bio3 | 2.1 | Gst | 2.2 |
| Bio7 | 1.8 | Bio7 | 1.1 |
| Bio4 | 1.8 | Gsp | ≈ 0 |
| Gsp | 1.5 |  |  |

Out of 630 candidate models for the 2^nd^ Chelsa model (Model 2) built for future projection, 625 were statistically significant (pROC test p < 0.05). Out of these, 201 met the 5% omission rate criteria (≈4.5%), and only 2 of them met the AICc criteria: a PH (product hinge) model with a regularization multiplier of 2 (ΔAICc ≈ 0), and a QPH (quadratic product hinge) model with a regularization multiplier of 2 (ΔAICc ≈ 1.27). Both of these final calibration models used the Set 1 variables created for the 2^nd^ model. Mean AUC values were 1.23 and 1.20 for the product hinge and quadratic product hinge models, respectively. The final evaluation with the independent test set showed omission rates 8.3% and 12.5% for the PH and QPH models, respectively, indicating that the PH model is better for predictions. For the 2^nd^ model, there were 11 predictors. The biggest contributors were Bio13 (precipitation amount of wettest month) with 31.3% followed by Bio14 (19.4% contribution) and Gdd0 (growth degree days above 0C) with a 13% contribution. Like Model 1, Bio6 was the 4^th^ important predictor for Model 2 with 8.2% contribution.

### Current and Future Predictions

Threshold values for creating binary maps were 0.409 and 0.483, respectively. Figure 1 shows the suitable areas for present conditions. According to the 1^st^ model, the current climatically suitable region for *Hyalomma marginatum* stretches out from Iberian Peninsula to Anatolia and the Caspian Sea. This region includes most of the Mediterranean Basin. This pattern mostly coincides with the reported locations of *H. marginatum*. The predictions of 2 models are mostly compatible with each other. Compared to the 1^st^ model, the prediction of the 2^nd^ model for the current distribution shows a wider suitable area in some regions, especially in North Africa and near Hungary. Also, the 1^st^ model shows wider suitable regions in Trans-Caucasus and Spain compared to 2^nd^ model. This might be due to the fact that the 2nd model is does not include parameters like cmi and rsds which are present in the 1^st^ model. On the other hand, the 2^nd^ model includes another important parameter, gdd0, which was eliminated from the 1^st^ model in the correlation and vif selection procedure.

**Figure 1.**
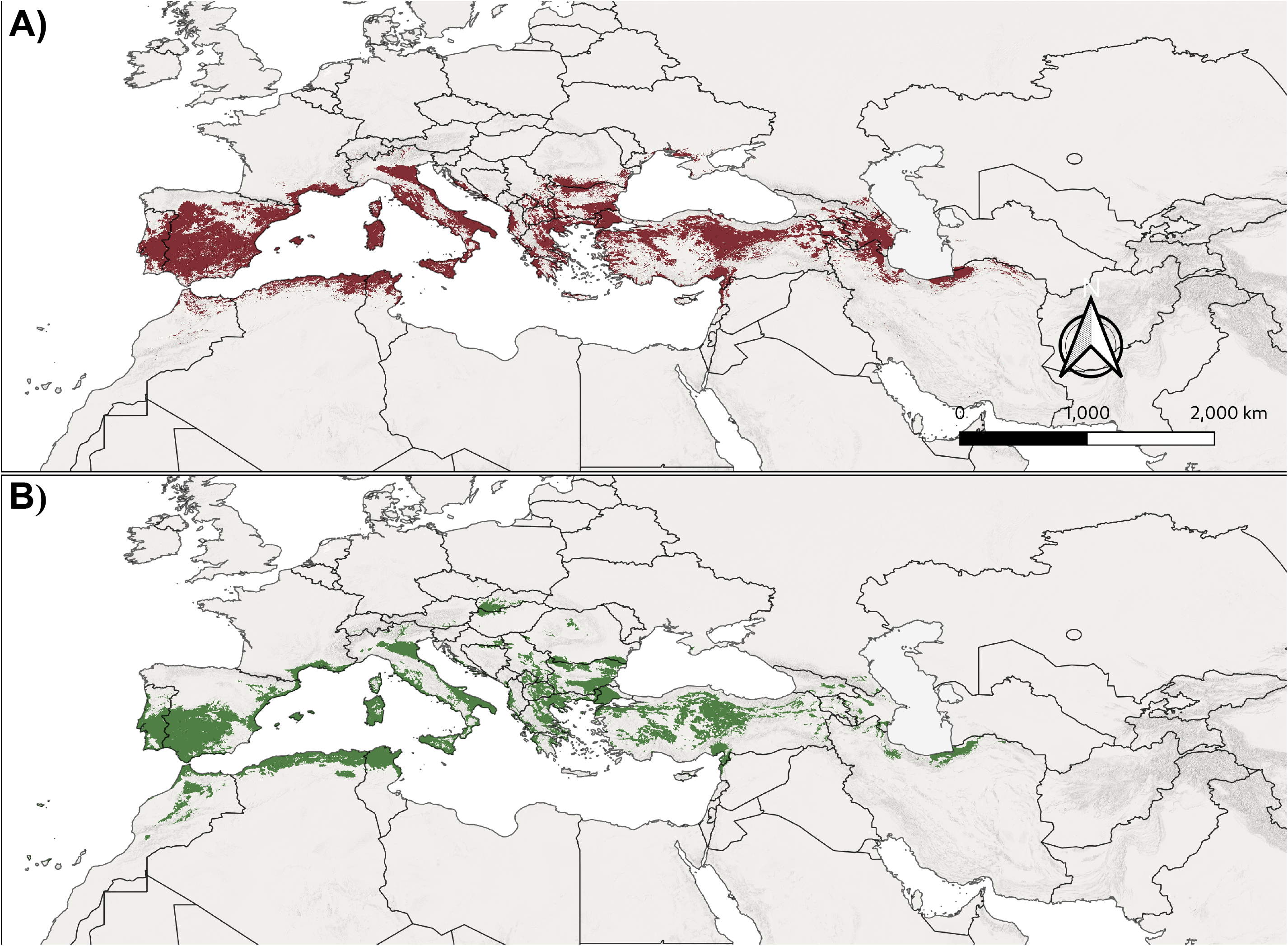
Maps of predicted suitable areas from the ENM results. **A)** green areas show the suitable regions under current conditions according to Model 1 **B)** red areas show the sutiable regions under currrent conditions for Model 2.

Figure 2 shows the predicted changes in suitable areas under GCM scenarios compared to present conditions. Future projections under ssp370 scenarios for 2011-2040 show a significant widening of the climatically suitable area. The direction of the expansion is towards north in Europe, westwards in Anatolia, and south in North Africa regions. Interestingly, the projections for the 2041-2070 period do not show such widening compared to the 2011-2040 period in Eastern Europe and Anatolia; for example, the patches of new barely suitable regions in Baltic regions and Russia are lost in the 2041-2070 scenarios. On top of that, there are significant declines in some regions in both ssp370 and ssp585 scenarios; these areas of suitability decline are especially distinct in Central Anatolia and some areas in Balkans and Central Europe. While both ssp370 and ssp585 show similar patterns in new areas for 2041-2070 period, ssp585 scenarios present a wider new region of low suitability in France and Eastern Germany.

**Figure 2.**
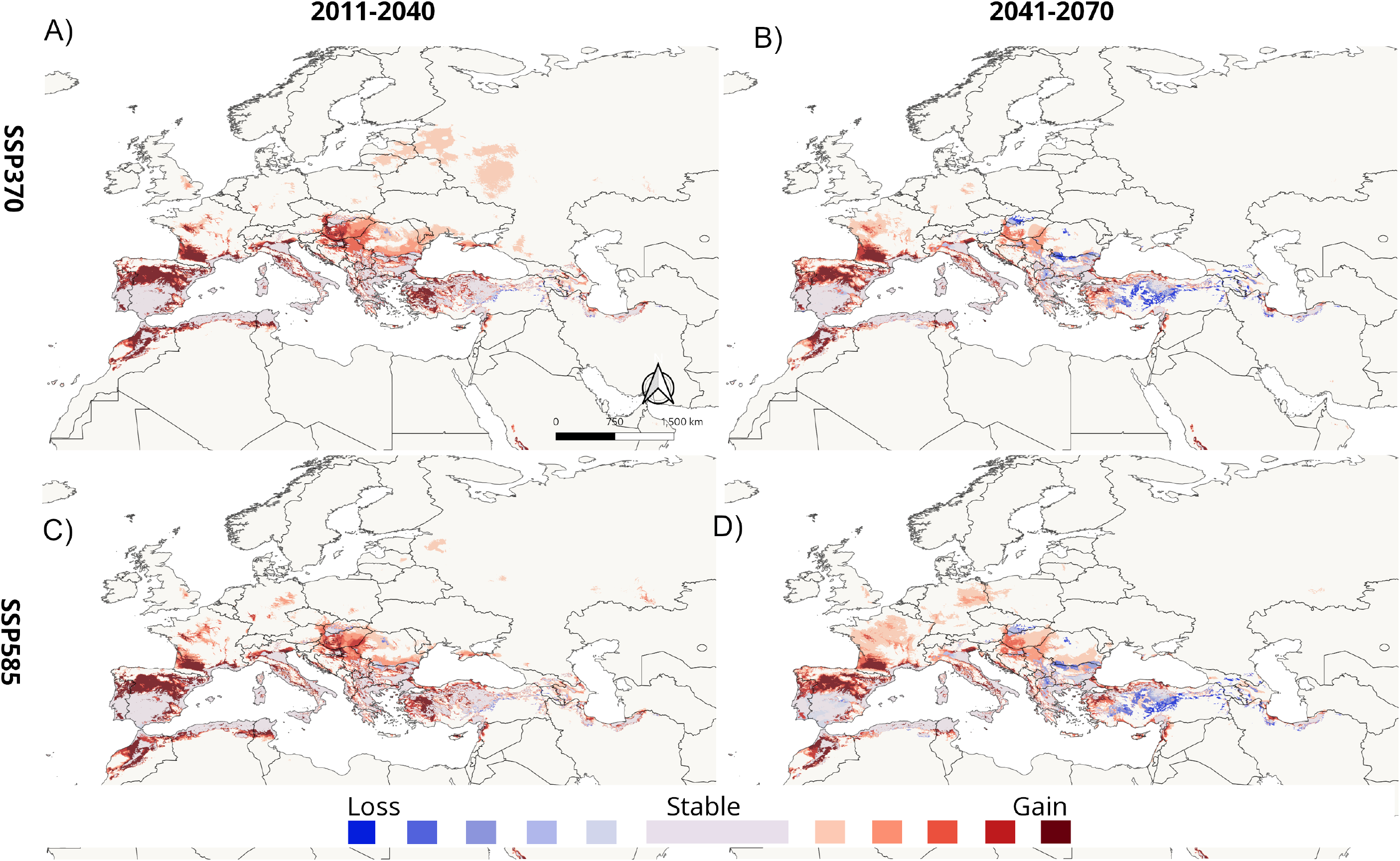
Maps of predicted suitable areas for future average of 5 GCM scenarios with differing degrees of loss and gain compared to current conditions for Model 2. **A)** for ssp370 in the 2011-2040 period **B)** for ssp370 in the 2041-2070 period **C)** for ssp585 in the 2011-2040 period **D)** for ssp585 in the 2041-2070 period.

## DISCUSSION

In this study, our aim was to predict the climatically suitable areas for present conditions and future scenarios using high-resolution updated bioclimatic variables and an updated list of locations for *Hyalomma marginatum*. The outputs from the first model show that the main climatic limiting factors for *H. marginatum* are climate moisture index and precipitation, followed by surface radiation and minimum temperatures. Relative humidity or moisture is crucial for ticks, as they are highly susceptible to cuticular water loss in their off-host state (Benoit & Denlinger, 2010; Estrada-Peña et al., 2012; Knülle & Rudolph, 1982; Leal et al., 2020; Requena-García et al., 2017). The importance of water content of air for *H. marginatum* has also been demonstrated in previous habitat suitability models (Estrada-Peña et al., 2011; Fernández-Ruiz & Estrada-Peña, 2021). An interesting new parameter in our model is surface radiation. Incoming solar radiation is, in fact, a critical parameter for small arthropods, which is generally overlooked (Barrett & O’Donnell, 2023; Battisti et al., 2013; Davis et al., 2005). Solar radiation might affect many different parameters related to ticks. Firstly, it is the main driver of microclimate temperature, and higher levels of solar radiation might increase tick abundance and questing activity (Del Fabbro et al., 2015; Kiewra et al., 2014). However, due to UVB radiation, very high levels of solar radiation might also have deleterious effects on small ectotherms like acari (Sakai et al., 2012; Sudo & Osakabe, 2015).

The predicted area for the current conditions in Model 1 is mostly compatible with previous habitat suitability models done for *H. marginatum* (Fernández-Ruiz & Estrada-Peña, 2021; Williams et al., 2015). Additionally, the model output shows potential suitable areas in Trans-Caucasus and Iran, especially in the Northern Iran near the coast of Caspian Sea, despite the dataset of locations we used for the models not including any locations from this region. Although no exact coordinates are presented in the literature, CCHF cases and also *H. marginatum* ticks carrying the virus have been reported from this area (Sedaghat et al., 2017; Shemshad et al., 2012; Sofizadeh et al., 2014; Telmadarraiy et al., 2015).

Model 2 output shows a larger climatically suitable area compared to the more restrictive predictions of the Model 1. Important parameters like moisture index (cmi) and surface radiation (rsds) were left out for this model, as they are not available for future scenarios. Moisture index is replaced by precipitation of the wettest period (Bio13) as the most important parameter in Model 2, which was in 6^th^ place in the previous model. Although net precipitation amount is not as strong as water deficit or moisture index as a predictor, this shows the importance of the water balance for climatic suitability. In addition, surface radiation is replaced by gdd0 (growth degree days above 0□), which was eliminated from Model 1 due to correlation threshold with other parameters. Fulfilling the required degree days is an important necessity for *H. marginatum* populations to establish in a new region (Estrada-Peña, 2023). It has been reported that an accumulated temperature of 3000-4000C is necessary for this species; a northern limit roughly coincides with 47C N (Estrada-Peña et al., 2011; Gillingham et al., 2023). With the increased temperature in recent decades, new reports of *H. marginatum* have also been increasing in Europe (Bah et al., 2022; Duscher et al., 2018; Hornok & Horváth, 2012; Lesiczka et al., 2022). Gillingham et al., (2023), who modelled the probability of survival and establishment of *H. marginatum* populations depending on cumulative degree days for the United Kingdom, forecasted increased risk in Southern UK in the future. Also, in our results, the prediction maps for the climatically suitable area for ssp370 and ssp585 scenarios point to a Northward expansion in the future. Depending on the models trained with trends in climate and tick records between 1970-2018, Fernández-Ruiz & Estrada-Peña, (2021), assumed that the suitable climatic area in Europe would expand while maintaining the original Mediterranean distribution. Additionally, Williams et al. (2015) also predicted a northward expansion under AR5 climate scenarios in the future. Another parameter that should be considered is minimum temperatures, which occupied 4^th^ place in both Model 1 and Model 2. In addition to the above-mentioned effect of temperature (degree days) on development time, overwintering survival is another important factor for *H. marginatum*, as most adults overwinter in the field (Valcárcel et al., 2020). An increase in the minimum temperatures in the field might increase winter survival and contribute to the ongoing and possible future shift in the northern limits of *H. marginatum* populations (Estrada-Peña et al., 2012). Our projections are compatible with these predictions, showing a very similar pattern of new suitable regions in North Spain, France, the Balkans and Western Anatolia. The projections for the 2040-2070 point to a continuation of this expansion; however, the projection also points to significant declines in some regions, mostly in Central Anatolia and the Balkans. This is most probably due to the decrease in precipitation, as climatic simulations predict that the impact of decreased precipitation will be much higher in the Eastern Mediterranean, especially in Turkey (Hoerling et al., 2012; Turkes et al., 2020). It was previously suggested that the northern limit for *H. marginatum* is thought to be mainly determined by temperature, while the southern limit is determined by precipitation and humidity (Gray et al., 2009), and expansion in the North and contraction in the South is an expected outcome in the future.

We also have to consider that in some regions, the change in suitability might be more complex than this pattern. With projected ongoing climate change, we can see an interchange of climatically suitable and unsuitable areas in models (Gillingham et al., 2023). For example, a previous model using RCP8.5 scenarios carried locally for Romania depicted an increase in suitable areas until 2050, followed by a decreasing suitability until 2070 (Domşa et al., 2016), which was also detected by our future predictions (Figure 2). This rather complex pattern in suitability might be due to the 2 main factors (temperature and precipitation) determining suitability. For instance, our present-day models show a highly suitable region for *H. marginatum* in the central and central-north parts of Turkey. Contrasting with the relatively more northward expansion in Europe, future projections indicate that after 2040, this suitable area might shift towards coastal areas where required precipitation would be available compared to the now more arid continental regions.

Assessing the potential distribution range of ticks is important to predict the risk of emergence and re-emergence of tick-borne diseases (Estrada-Peña, 2008; Zhao et al., 2021). In this sense, the interaction between ticks and climate is extremely important, as the survival and biological functions of ticks depend strongly on the external micro-abiotic environment (Gray et al., 2009). It should also be kept in mind that finding a suitable host is one of the most important drivers, which affects the distribution of tick species in an area. On the other hand, *Hyalomma marginatum* is a two-host species and uses a wide range of vertebrates at different stages of its life cycle; this complex behaviour of this species is another reason for the difficulties in predicting the future distributions of these vectors (Apanaskevich & Horak, 2008; Bonnet et al., 2022; Valcárcel et al., 2020).

Ecological niche models are generally more deterministic models that build the representative fundamental niche using presently available abiotic parameters and locations, then project this assumed niche space to assumed future abiotic parameters data calculated by predictive global circulation simulations (Escobar, 2020; Sillero et al., 2021). Thus, our projections are for predicting the climatically suitable areas in the predicted future under assumed scenarios. However, these kinds of models are among the most useful tools presently available for estimating the potential risk of vector-borne diseases and deciding the necessary precautions (Medlock & Leach, 2015). With the ongoing human-induced changes in the biotic and abiotic environment we have been observing a significant increase in dispersal and abundance of *H. marginatum* and also an increase in the transmission of pathogens carried by *H. marginatum*. Future projections indicate a significant climb in potential risk due to these factors, so it is important to build both statistical and process-based mechanistic models with new data. We also strongly suggest publishing coordinates, as we have seen that there are many studies which report species presence without any coordinates that can help future coordination between researchers and provide more reliable models.

## Author contributions

Conceptualization, O.H., and A.C.K.; methodology, A.C.K. and C.E.; data curation, O.H., formal analysis, A.C.K. and C.E.; interpretation; A.C.K. and C.E.; writing-original draft preparation, A.C.K.; writing-review and editing O.H., A.C.K. and C.E.; visualization, C.E.

## Acknowledgments

The numerical calculations reported in this paper were partially performed at TUBITAK ULAKBIM, High Performance and Grid Computing Center (TRUBA resources).

